# The capsular polysaccharide obstructs wall teichoic acid functions in *Staphylococcus aureus*

**DOI:** 10.1101/2023.07.26.550747

**Authors:** Esther Lehmann, Rob van Dalen, Lisa Gritsch, Christoph Slavetinsky, Natalya Korn, Carina Rohmer, Daniela Krause, Andreas Peschel, Christopher Weidenmaier, Christiane Wolz

## Abstract

**Background:** The cell envelope of *Staphylococcus aureus* contains two major secondary cell wall glycopolymers: capsular polysaccharide (CP) and wall teichoic acid (WTA). Both the CP and the WTA are attached to the cell wall and play distinct roles in *S. aureus* colonization, pathogenesis, and bacterial evasion of host immune defenses.

**Objective:** We aimed to investigate whether CP interferes with WTA-mediated properties.

**Methods:** Strains with natural heterogeneous expression of CP, strains with homogeneous high CP expression and CP-deficient strains were compared to WTA deficient controls regarding WTA dependent phage binding, cell adhesion, IgG deposition, and virulence *in vivo*.

**Results:** WTA-mediated phage adsorption, specific antibody deposition and cell adhesion were negatively correlated with CP expression. WTA, but not CP, enhanced the bacterial burden in a mouse abscess model, while CP overexpression resulted in intermediate virulence *in vivo*.

**Conclusions:** CP protects the bacteria from WTA-dependent opsonization and phage binding. This protection comes at the cost of diminished adhesion to host cells. The highly complex regulation and mostly heterogeneous expression of CP has probably evolved to ensure the survival and optimal physiological adaptation of the bacterial population as a whole.

## Introduction

The bacterial cell envelope is a multilayered structure that mediates specific interactions with the changing environments encountered during infection or colonization. The surface of *Staphylococcus aureus* is highly glycosylated and is mainly composed of peptidoglycan (PG), cell wall-anchored (often glycosylated) proteins and secondary cell wall glycopolymers such as wall teichoic acid (WTA) and capsular polysaccharide (CP). Both WTA and CP are covalently linked to the C-6 hydroxyl of MurNAc of PG [1, 2]. *S. aureus* WTA is composed of 30-50 ribitol-phosphate (RboP) repeating units that are modified with D-alanine and GlcNAc [3, 4]. WTA has multiple functions in cell wall turnover, cellular physiology, phage interaction, host cell adhesion, antibiotic resistance, and immune interaction [4-8]. High quantities of WTA, as found in the clinically highly relevant, methicillin-resistant USA300 lineage, have been linked to increased virulence [9]. In addition, as the primary staphylococcal phage receptor, WTA is also of critical importance for *S. aureus* genetic plasticity, as horizontal gene transfer in *S. aureus* is driven largely by phage transduction. The GlcNAc modifications of WTA are important for interaction with some, but not all, *S. aureus*-targeting phages [7, 10].

CP serotypes 5 and 8 are the two main CP serotypes produced by *S. aureus* [6, 11]. The proteins involved in CP5 or CP8 biosynthesis, *O*-acetylation, transport and regulation are encoded by allelic *cap5A-P* or *cap8A-P* operons, respectively. The structures of CP5 and CP8 are very similar, both consisting of trisaccharide repeating units of D-*N*-acetyl mannosaminuronic acid, L-*N*-acetyl fucosamine, and D-*N*-acetyl fucosamine. The difference between the serotypes lies in the linkages between the sugars and the site of *O*-acetylation of the mannosaminuronic acid residues [12-14]. CP expression depends on the local conditions encountered during particular types of infections [6, 11, 15]. CP enhanced virulence in murine models of bacteremia [16, 17], septic arthritis [18], and surgical wound infection [19]. Furthermore, loss of CP expression has been reported in *S. aureus* isolates from chronic infections such as osteomyelitis [20], mastitis [15], cystic fibrosis [21] or atopic dermatitis [22] indicating that loss of CP expression is of advantage under these conditions. Moreover, acapsular *S. aureus* derived from chronic infections can regain capsule expression after passage through the bloodstream in a bacteremia mouse model [23]. In contrast, in other animal models CP expression has been linked to reduced virulence [24-26].

Interestingly, the highly virulent USA300 lineage is CP deficient due to conserved mutations in the *cap5* locus [27]. The seemingly contradictory role of CP in either enhancing or diminishing virulence is reflected by the highly heterogeneous expression of CP. At the population level, CP has a unique expression pattern tightly linked to the growth phase of the bacterium [6]. *Cap* expression is repressed during exponential growth, starts heterogeneously in the late exponential phase and reaches its maximum during the stationary phase while still featuring highly heterogeneous expression at the population level. The heterogeneous synthesis of CP entails a highly complex regulation of *cap* expression and the interplay between several regulatory circuits and environmental conditions. The alternative sigma factor B (SigB) and the extended region upstream of the SigB consensus sequence within the *cap* promotor (P_cap_) region controls *capA-P* expression. Mutation of the upstream region and modification of the SigB binding motif result in strains with increased and more homogenous CP expression [6].

We hypothesize that the highly heterogeneous pattern of CP expression is beneficial for the *S. aureus* population. The CP-positive cells may allow immune and phage evasion, whereas the CP-negative ones may favor adhesion since CP likely masks the function of cell-wall attached adhesins such clumping factor A [28, 29]. For the first time, we analyzed the functional impact of CP expression on another cell wall polysaccharide and show that CP obstructs major functions of WTA with regard to adhesion, phage absorption and IgG deposition.

## Methods

### Bacterial strains, phages, and growth conditions

All bacterial strains and phages are listed in the supplemental information in Table S1 and Table S2 respectively. Bacteria were cultivated overnight in TSB (tryptic soy broth, Oxoid/Difco Laboratories) (37°C, 200 rpm). Resistant bacteria were grown with appropriate antibiotics (ampicillin [10 µg/ml], erythromycin [2.5 µg/ml], chloramphenicol [10 µg/ml], kanamycin [50 μg/ml]) during overnight culture.

### Strain construction

The plasmids and oligonucleotides used in this study are listed the supplemental information in Table S3 and Table S4, respectively. A markerless Δ*tagO* mutant was generated in *S. aureus* USA300 JE2 by Gibson assembly of the two flanking regions of *tagO*, amplified using the primer pairs tagOGibMutfor/tagOGibMutrev and tagOlinkMutfor/tagOlinkMutrev, into the mutagenesis vector pBASE6, followed by mutagenesis as previously described [30]. An internal deletion of *int* (SAOUHSC_02239) of phi13kan was generated using the mutagenesis vector pIMAY [31]. Briefly, the right and left *int* flanking regions were amplified using the primer pairs pCG849i1gibfor/pCG849i1gibrev and pCG849i2gibfor/pCG849i2gibrev and linked via PCR. The fragment was cloned into the shuttle vector pIMAY by Gibson assembly, cloned into *E. coli* DC10B and transferred into USA300cФ13kan by electroporation. Mutagenesis was performed as previously described [31]. Deletion of the N-terminal part of *int* in USA300cФ13kan-Δ*int* was verified via PCR. To obtain Ф13kan-Δ*int* (which is unable to excise from the bacterial chromosome), we complemented USA300cФ13kan-Δ*int* with its native integrase using plasmid pCG32 [32]. pCG32 was transduced from RN4220 into USA300cФ13kan-Δ*int* using Ф11. Ф13kan-Δ*int* was induced from USA300cФ13kan-Δ*int*, pCG32 with mitomycin C (500 ng/ml). All strains were verified on blood agar plates for Agr activity.

### WTA isolation and purification

WTA isolation was modified from Peschel *et al*. 1999 [33]. Briefly, *S. aureus* was grown overnight in B-Medium (5 g/L yeast extract (Ohly Kat, Deutsches Hefewerk), 10 g/L soy peptone (A3SC, Organo Technie), 5 g/L NaCl, 1 g/L glucose, 0.8 g/L K_2_HPO_4_) with an additional 0.2% glucose. Bacteria were disrupted by mechanical forces using a FastPrep-24 (MP Biomedicals) and glass beads (Sigma, 425-600 µm). After sedimentation by centrifugation, the lysate was digested with DNase/RNase (Sigma) and treated with 2% SDS. WTA was released from the peptidoglycan by treatment with 5% trichloroacetic acid and separated by centrifugation. The amounts of phosphorus in the WTA samples were determined as described previously in Chen *et al*. [34] against a phosphate standard solution (Sigma, P3869). The pellet was dried via vacuum centrifugation (Speedvac), and the amount of peptidoglycan (PG) was determined by pellet weight. The amount of WTA/PG was displayed as [nmol _WTA_/mg _PG_].

### PFU determination via plaque assay

Phage titer was determined by the agar overlay method as described previously [35] using *S. aureus* strains RN4220 (for Φ11 and ΦK) or LS1 (for Φ13k∆*int*) as indicator strains. Briefly, indicator strains were grown to OD_600_ = 0.1 in TSB. 100 µl of bacterial cultures were mixed with 3 ml liquid soft agar (TSA + 0.75% agar) and poured on an NB2 plate (Nutrient Broth No. 2 (Oxoid), 0.75% agar, 2.72 mM CaCl_2,_ pH 7.5). After solidification, 10 µl of diluted, sterile filtered phage lysates (obtained using a 0.45 µm membrane filter (Labsolute)) were spotted on the lawn and plaque forming units (PFU) determined.

### Phage adsorption assay

Phage adsorption assays were performed at an multiplicity of infection (MOI) of 0.01 as described previously [10]. Briefly, 10^8^ bacterial CFU from an overnight culture were incubated with 10^6^ phages in a total volume of 1 ml of TSB with 2.72 mM CaCl_2_ for 10 min at room temperature under non-shaking conditions. Bacteria were pelleted (5000 x g, 5 min), and the supernatant was sterile filtered using a 0.45 µm membrane filter and used for PFU determination.

### Antibody deposition

Antibody deposition to *S. aureus* WTA was performed analogously to Hendriks *et al*. [36] employing an anti-α-1,4-GlcNAc-WTA Fab (clone 4461) [37] to avoid epitope-independent interaction with staphylococcal protein A. In brief, bacterial cultures were adjusted to 1.25*10^6^ cells in 1x PBS (phosphate-buffered saline) with 0.1% BSA (bovine serum albumin; Sigma A7906) and incubated for 30 min at 4°C with 4461-Fab at 10 µg/ml - 0.1 µg/ml in 3-fold serial dilution steps. After washing with PBS/ 0.1% BSA, bacteria were incubated with 1:1000 diluted PE-conjugated F(ab’)2-anti-human-kappa-light-chain (Invitrogen, PA1-74408) in PBS/ 0.1% BSA (20 min, 4°C). Unbound antibodies were removed, and bacteria were fixed in 1% paraformaldehyde (PFA) in PBS for 15 min. The fixative was removed, and bacteria were resuspended in a total volume of 150 µl PBS for flow cytometric analysis. Flow cytometry data were acquired on a FACSFortessa (BD) using FACSDiva (BD). Per sample, 10,000 events were measured within the set gate. All data were analyzed using FlowJo 10 (FlowJo, LLC).

### FITC labeling of bacteria

10 ml bacterial overnight cultures were washed three times, resuspended in 1x PBS and labeled with 0.1 mg/ml fluorescein isothiocyanate (FITC) (Sigma) at 4°C for 1 h. Labeled bacteria were washed three times with 1x PBS and resuspended in DMEM-F12 (Gibco). Labeled bacteria were stored at -20°C. Bacterial cell numbers were determined using a Neubauer counting chamber.

### Cell culture and adhesion assays

Human umbilical vein endothelial cells (HUVECs) from pooled donors (Promocell) were cultured in endothelial cell growth medium (Promocell) and used up to passage number six (see supplemental information Table S5). Cell culture and incubation during the experiments were performed at 37°C under 5% CO2 and 95% rH. During experimental procedures, HUVECs were kept in DMEM-F12 (Gibco).

Adhesion assays were performed under static conditions with HUVECs and FITC-labeled bacteria. Briefly, HUVECs were seeded at a density of 1 to 3×10^6^/channel in channel slides (ibidi µ-Slide VI 0.4) and incubated to attach overnight at 37°C, 5% CO_2_ and 95% rH. Prior to infection, HUVECs were washed once with prewarmed DMEM-F12 (Gibco). For the USA300 JE2 strain panel, pilot experiments confirmed an optimal window of resolution of 1 h at MOI 10 (data not shown). For the Newman strain panel, we confirmed an optimal window of resolution of 2 h at MOI 10 (data not shown). After infection, the channels were washed twice with prewarmed DMEM-F12 and once with prewarmed 1x PBS (Gibco). Cells were fixed with 4% PFA in 1x PBS (Gibco) for 20 min at RT and permeabilized with 0.1 % Triton X-100 in 1x PBS for 10 min at RT. DNA was stained with DAPI (Thermo Fisher, [1 µg/ml]) for 20 min at RT, and the actin cytoskeleton was stained with phalloidin-TRITC (Thermo Fisher, [2.5 µg/ml]) for 30 min at RT. The number of bacteria adhering to HUVECs was determined by confocal microscopy on a Zeiss LSM710 NLO.

### Abscess model

The murine abscess model procedures were altered from Wanner *et al*. [9]. Briefly, eight-week-old female C57BL/6JRccHsd mice (Envigo) were housed with up to five animals per cage in individually ventilated cages (IVCs). Bacterial suspensions from overnight cultures were adjusted to a CFU of 10^5^ in 100 µl and mixed with an equal volume of dextran beads (Cytodex-1 microcarriers; Sigma) to obtain a final concentration of 0.5% cytodex-1 beads per 200 µl. Mice were anesthetized using an isoflurane-oxygen mixture (cp-pharma) and injected subcutaneously in each flank with 0.2 ml of the *S. aureus*-bead suspensions. Mice were tightly monitored and euthanized after two days of bacterial challenge. The abscesses were excised and homogenized in 1 ml 1x PBS containing protease inhibitor (Roche) according to the manufacturer’s instructions. Serial dilutions of the homogenates were plated in technical duplicates on TSA plates. The results were analyzed as the log CFU of bacteria/abscess.

### Statistical analysis

For statistical analysis, we used GraphPad Prism (GraphPad Prism Software 9.5.0) and appropriate statistical methods as indicated in the figure legends. P < 0.05 was considered significant. All statistical analyses were based on independent biological replicates.

## Results

### WTA abundance is unaffected by CP production

Although WTA and CP are synthesized by dedicated biosynthesis enzymes encoded by separate gene clusters, these biosynthesis pathways are entangled. Both WTA and CP rely on the universal UDP-GlcNAc building blocks, the universal lipid II (UDP) carrier and mutual enzymes from the LPC family for attachment to the PG. We therefore analyzed whether the WTA content is affected by the CP synthesis. We used a set of *S. aureus* mutants with varying CP synthesis capacities: *S. aureus* strain USA300 JE2 is naturally CP-deficient as a result of three point-mutations within *capD, capE* and the *cap* SigB consensus motif [27]. These three mutations have been cured in USA300 JE2 Cap, resulting in a mutant with canonical heterogeneous CP expression in the stationary phase [6]. USA300 JE2 Cap-high is a derivative of Cap that homogeneously expresses CP in the stationary phase, which was achieved by deletion of the repressive upstream region and mutation of the SigB binding motif of the *cap* promotor region.

We measured the total WTA abundance per dry cell wall weight in the two strains with the largest difference in CP production, the CP-deficient wt and Cap-high (figure 1). WTA was extracted from stationary phase cultures to ensure high CP expression. The WTA abundance did not differ significantly between these two strains, indicating that CP deposition does not interfere with WTA linkage to the cell wall.

**Figure 1.**
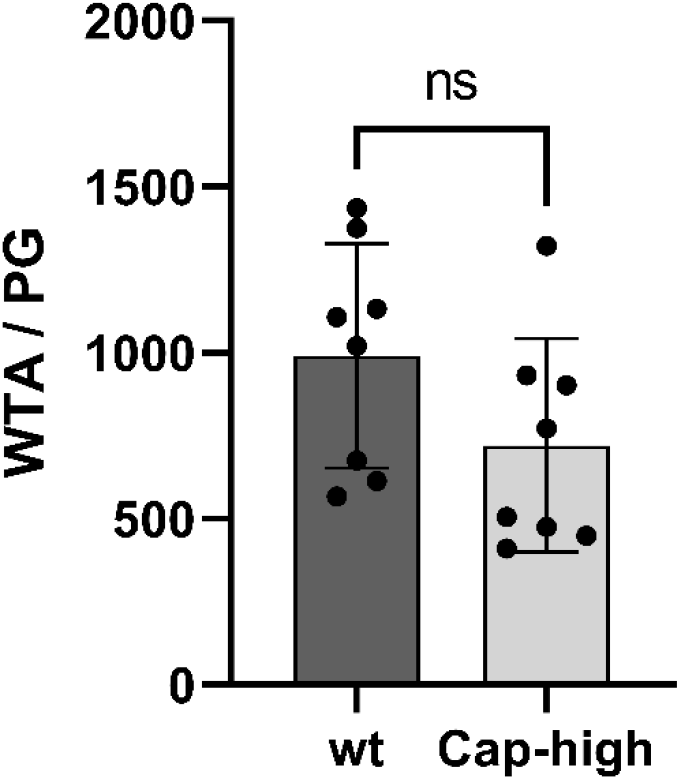
Impact of CP expression on WTA abundance. WTA of USA300 JE2 (wt) and Cap-high was isolated and quantified from stationary phase cultures. The WTA abundance was approximated by the phosphate concentration in nmol per mg peptidoglycan (PG). Data are presented as the mean and SD of eight independent biological replicates. Statistical analysis was performed by unpaired t test. No statistically significant (ns) differences were found.

### CP expression reduces phage absorption

We assessed whether CP affects WTA-mediated phage absorption. Strains with varying degrees of CP, as well as the WTA-deficient USA300 JE2 Δ*tagO* mutant, were incubated with representative phages: Φ11 (siphovirus, shown to recognize α-1,4-GlcNAc-, β-1,4-GlcNAc-and partially β-1,3-GlcNAc-RboP-WTA [7, 10]), ΦK (myovirus, shown to recognize RboP-WTA [10]) and Φ13 (siphovirus, unknown phage receptor). Since Φ13 phages produce turbid plaques that are difficult to enumerate, we partially deleted the phage integrase gene *int*, resulting in a phage derivative Φ13 Δ*int* that produces clear plaques. After adsorption, the number of unbound phages was determined in a double agar overlay plaque assay. Low numbers of phages in the supernatants indicate a high rate of phage adsorption (figure 2).

**Figure 2.**
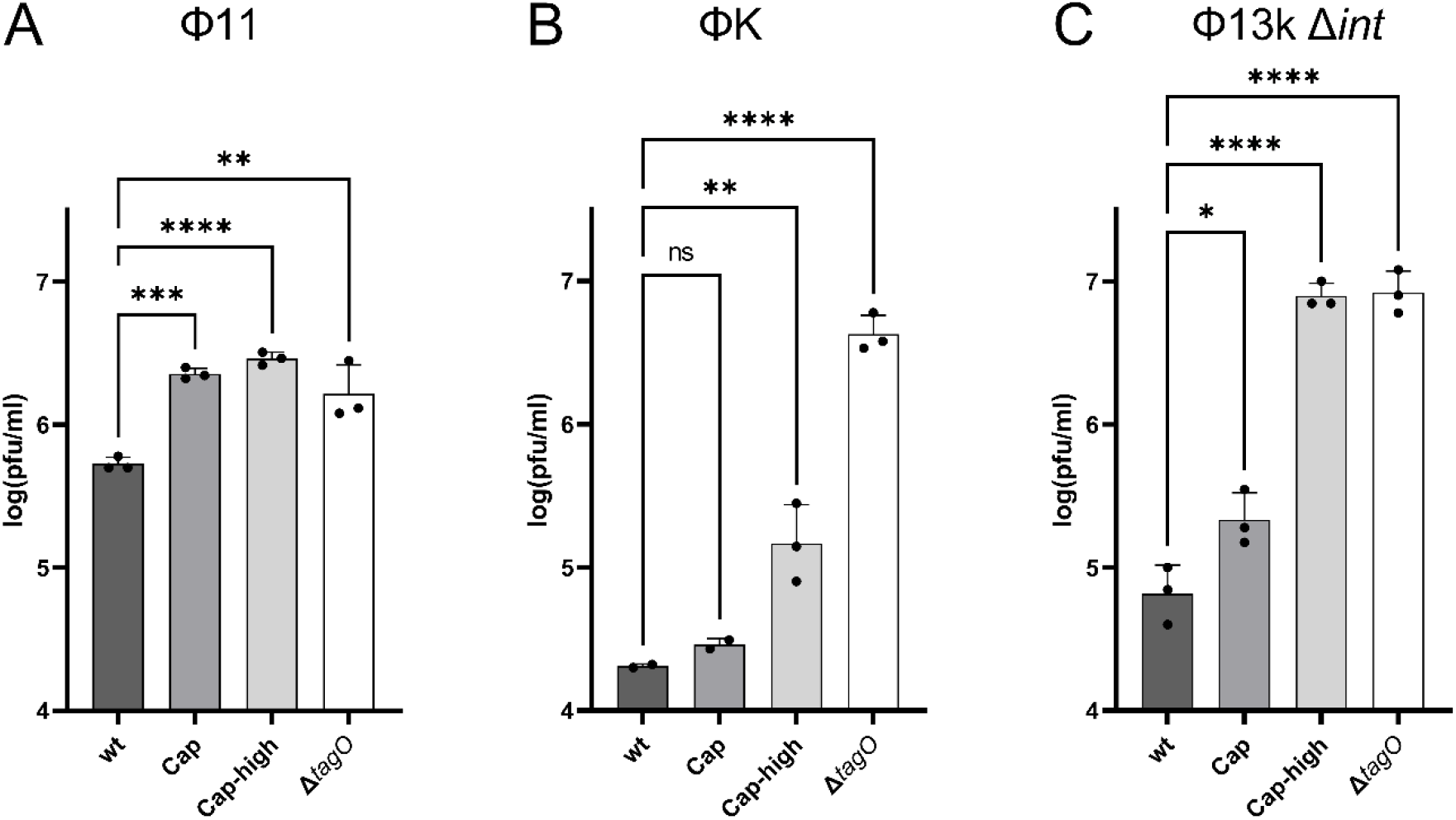
CP expression reduces phage absorption. Plaque-forming units (PFU) after adsorption of phages to S. aureus cell wall mutants (MOI 0.01, 10 min). USA300 JE2 wt: CP-deficient; Cap: heterogeneous CP expression; Cap-high: homogeneous high CP expression; ΔtagO: WTA deficient. Higher phage adsorption is indicated by reduced numbers of unbound phages. **A**: Adsorption experiment with Φ11. **B**: Adsorption experiment with ΦK. **C**: Adsorption experiment with Φ13k∆*int*. Represented are log-transformed data with mean and SD of three independent experiments. Statistical analysis was performed by one-way analysis of variance (ANOVA) with Dunnett’s test for multiple comparisons. Significant differences vs. the wt are indicated by one (P<0.05), two (P<0.01), three (p<0.001), or four (P<0.0001) asterisks (*).

No absorption was detected in the ∆*tagO* mutant, consistent with the dominant role of WTA as a phage receptor. Phage adsorption was negatively correlated with the CP expression level. For all three phages, the highest adsorption was observed for the CP-negative wild type, and adsorption to the CP-positive strains Cap and Cap-high was significantly decreased. For both Φ11 and Φ13k ∆*int*, absorption to Cap-high was as low as that to the WTA-deficient ∆*tagO* mutant (figure 2A and 2C), indicating that high CP expression efficiently blocks access of all three phages to their WTA receptor.

### CP expression reduces the binding of WTA-specific antibodies

An estimated 70% of *S. aureus*-reactive IgG is directed toward WTA GlcNAc [38]. We analyzed whether CP impedes access of these antibodies to WTA. To avoid the interaction of antibodies with the *S. aureus* antibody-binding protein SpA, we used anti-α-1,4-GlcNAc-F(ab’) fragments [39] in a flow cytometry-based immunofluorescence staining experiment. Antibody deposition on USA300 increased in a dose-dependent manner, reaching saturation at 3 µg/ml. Deposition was clearly impaired in the ∆*tagO* mutant and in the Cap-high strain (figure 3A).

**Figure 3.**
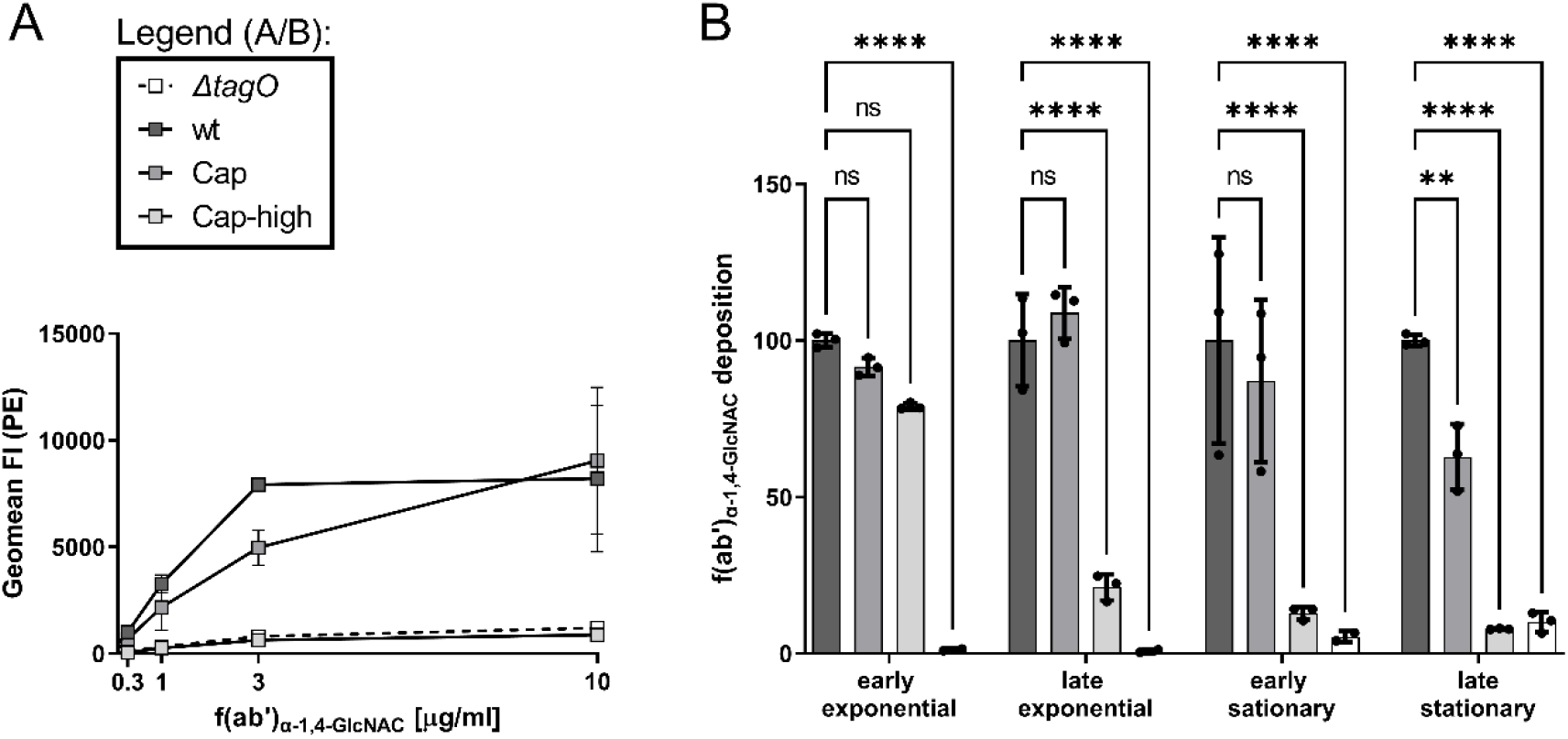
CP expression reduces the binding of WTA-specific antibodies. Binding of human α-1,4-GlcNAc-F(ab’) to S. aureus cell wall mutants (USA300 JE2 wt, Cap, Cap-high and ∆*tagO*), measured by flow cytometry using PE-conjugated human kappa light chain as the secondary antibody. **A**: Dose‒ response curve with 0 µg/ml, 0.3 µg/ml, 1 µg/ml, 3 µg/ml and 10 µg/ml α-1,4-GlcNAc-F(ab’). Data are displayed as the geometrical mean of the PE fluorescence intensity. **B**: α-1,4-GlcNAc-F(ab’) deposition at 3 µg/ml in bacteria form different growth phases as indicated. Shared figure legend in A. Data are displayed as the geometrical mean of the PE fluorescent intensity normalized to the wild type in the respective growth phase. The mean and SD of three independent experiments are shown. Statistical analysis was performed by two-way ANOVA with Dunnett’s test for multiple comparisons. Significant differences vs. the wild type are indicated by one (P<0.05), two (P<0.01), three (p<0.001), or four (P<0.0001) asterisks (*).

CP expression is highly growth-phase dependent, and bacteria from the exponential phase are mostly CP negative. Accordingly, CP-dependent inhibition of antibody deposition correlated with the growth phase. A significant reduction in WTA staining was observed only in bacteria from late exponential or stationary growth phase cultures (figure 3B). These results indicate that CP production limits the accessibility of antibodies to WTA, the dominant antibody target of *S. aureus*.

### CP expression reduces WTA-dependent adhesion to endothelial cells

CP expression in *S. aureus* strain Newman was shown to inhibit bacterial binding to endothelial cells, indicating that CP likely masks the major adhesins [28]. In strain Newman, *fnbA* and *fnbB* coding for fibronectin binding protein A and B are truncated [40]. As in USA300, WTA might function as a major adhesin in this strain background.

Endothelial cell adhesion assays using Δ*tagO* and the Sortase A-deficient mutant *srtA::erm* of strain Newman confirmed that endothelial cell adhesion is mainly mediated via WTA (figure 4). Moreover, Newman Cap-high showed significantly impaired adhesion to endothelial cells. In the *fnbAB* positive strain USA300, adhesion was also predominantly WTA dependent and negatively correlated with CP expression (Figure 4 B). These observations support that WTA is the main adhesin for endothelial cells and that CP significantly impedes this WTA-mediated binding.

**Figure 4.**
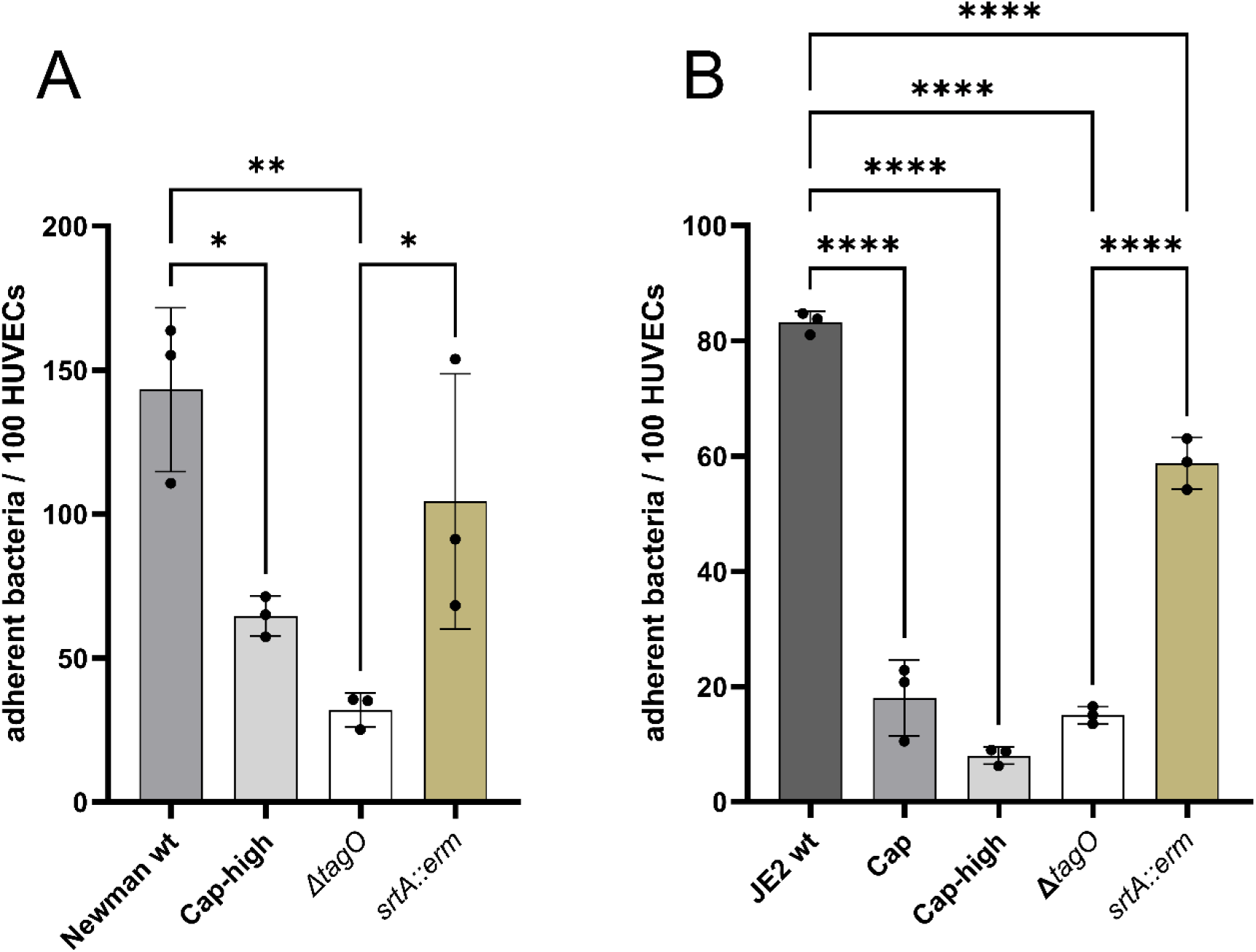
CP expression reduces WTA-dependent adhesion to endothelial cells. Adhesion of FITC-labeled bacteria to HUVECs. **A**: Newman and the isogenic derivatives Cap-high, ∆*tagO* and *srtA*::*erm* at MOI 10, 2 h. **B**: in the USA300 JE2 background wt, Cap, Cap-high, Δ*tagO* and *srtA::erm* at MOI 10, 1 h. Adherent bacteria were microscopically evaluated in five random fields of view per replicate. Data are displayed as the number of adherent bacteria per 100 HUVECs. The mean and SD of three independent experiments are shown. Statistical analysis was performed by one-way ANOVA with Tukey’s test for multiple comparisons. Significant differences are indicated by one (P<0.05), two (P<0.01), three (p<0.001), or four (P<0.0001) asterisks (*).

### CP expression reduces abscess formation

Previously, high WTA expression was shown to increase the bacterial burden in a mouse skin abscess model [9]. We hypothesized that CP production may affect this WTA-mediated phenotype. We therefore used an identical *in vivo* model and induced abscess formation in the flanks of C57BL/6J mice by injection of *S. aureus* mixed with Cytodex beads (figure 5) using the USA300 JE2 panel.

**Figure 5.**
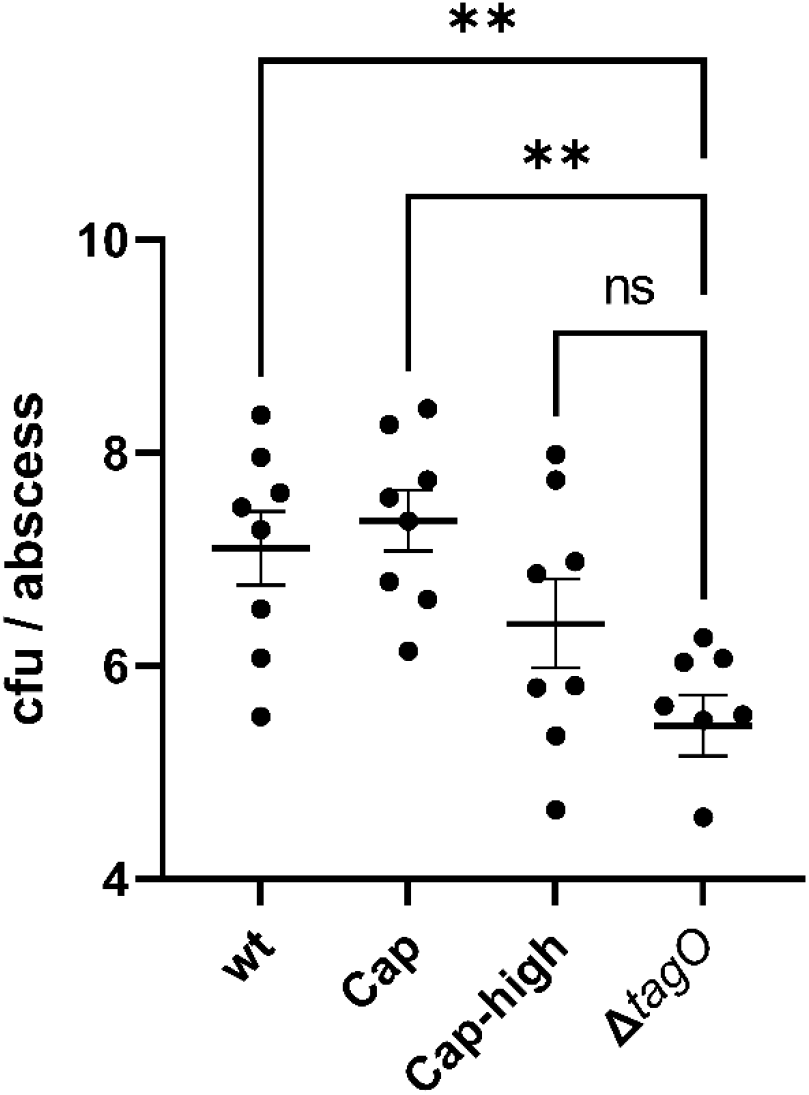
Impact of CP on WTA-mediated virulence in skin abscess formation. Abscess formation was induced by 10^5^ colony-forming units (CFU) of USA300 JE2 wt, Cap, Cap-high and Δ*tagO*. Bacteria were mixed with Cytodex beads and injected subcutaneously into the flanks of C57BL/6J mice (two abscesses per mouse). After 48 h, the mice were euthanized, the abscesses were excised and homogenized, and the bacterial burden was determined by dilution plating. Data are represented as a dot-blot with mean and SEM of the log-transformed CFU count. Each dot represents a single abscess. Statistical analysis was performed by one-way ANOVA with Dunnett’s test for multiple comparisons. Significant differences are indicated by one (P<0.05) or two (P<0.01) asterisks (*).

After 48 h, we observed a significantly reduced bacterial burden when mice were inoculated with Δ*tagO* compared to the wt. This confirms the pivotal role of WTA in abscess formation. The bacterial burden of Cap did not differ from that of the wt. High expression of CP resulted in an intermediate phenotype with a tendency toward lower bacterial burden compared to the wt and no significant difference to the WTA-deficient Δ*tagO* mutant. These results indicate that in this infection model, WTA, but not CP, is important for bacterial virulence and suggest a limiting effect of CP with regard to WTA-mediated virulence in skin abscess formation.

## Discussion

The central role of *S. aureus* secondary cell wall glycopolymers in virulence and host adaptation is widely discussed [4, 6, 41]. In this study, we show that CP impedes WTA functionality, such as phage absorption, cell adhesion and antibody deposition.

*S. aureus* WTA was previously shown to mediate adhesion to epithelial cells [42], Langerhans cells [43] and endothelial cells [44] (manuscript in submission), via recognition of host receptors SREC-1, langerin and LOX-1 respectively. Here, we confirmed that WTA is the major adhesin mediating the binding of *S. aureus* to endothelial cells. This could be shown for the *fnbAB-*deficient strain Newman as well as for the CP-negative strain USA300 JE2. In both strain backgrounds, adhesion was highly dependent on WTA. A *srtA* mutant which lacks cell-wall anchored proteaceous adhesins was only slightly diminished in its binding capacity to endothelial cells. Interestingly, multiple emerging *S. epidermidis* lineages express *S. aureus*-type WTA, which coincides with increased endothelial binding [45]. Overexpression of CP significantly decreased adhesion in Newman and USA300 JE2, confirming that CP masks WTA accessibility independent of the genetic background. The natural mutations within the *cap* operon as present in the USA300 lineage enhance the binding to endothelial cells (Figure 4A) in line with previous results using a *cap* deficient mutant of strain Newman [28]. CP likely also inhibits WTA dependent binding to other cells such as epithelial cells for which WTA also function as major adhesin [42].

WTA is the major phage receptor in *S. aureus* and other bacillota [46]. However, in many other species, including *Lactococcus lactis*, CP functions as the predominant phage receptor [47]. Here, we show that CP in *S. aureus* modulates phage absorption by impeding the binding of phages to WTA. Thereby, CP expression might hamper the efficiency of phage therapies. A similar observation was made previously in *Bacillus anthracis*, where shielding of phage absorption by CP was also demonstrated [48]. Previous observations regarding the role of CP in phage absorption in *S. aureus* were often inconclusive, and only the “macrocapsule” encoded by a different gene cluster were shown to interfere with phage absorption [49]. The results concerning the more prevalent CP5 and CP8 microcapsule were less clear [50, 51]. This might be explained by the observation that CP expression only occurs in a subpopulation of bacteria, specifically in the stationary growth phase [52]. Thus, phage inhibitory effects of CP in exponentially growing bacteria might be undetectable.

It was shown previously that when bacteria were grown under conditions of optimal CP synthesis, the CP protected from opsonophagocytic killing in the presence of either complement or antibodies [16, 53]. WTA is a major antigen of *S. aureus* as approximately 70% of all natural antibodies directed against *S. aureus* recognize WTA [38]. These antibodies are able to induce complement activation (Jung et al. 2012) and phagocytosis by neutrophils (Lehar et al. 2015; Dalen et al. 2019). Here, we demonstrate that enhanced CP production blocks opsonization by specific monoclonal anti-WTA-F(ab’) fragments [37]. To which extend CP contributes to immune-evasion during human infections remains unclear.

Interestingly, in early serotyping assays, agglutination of *S. aureus* in the presence of teichoic acid antiserum was used as a negative control to identify CP-deficient *S. aureus* isolates [12]. Until now, this observation has never been evaluated in the context of CP-WTA interactions.

The USA300 lineage is known for its high prevalence in skin and soft tissue infections, which has partially been ascribed to its WTA^high^ phenotype [9]. Interestingly, CP expression in *S. aureus* Reynolds was also shown to promote subcutaneous abscess formation [54]. Here, we confirmed the role of WTA for skin abscess formation, whereas CP expression did not promote abscess formation. Rather, the USA300-high strain displayed a phenotype more similar to the significantly attenuated WTA-deficient mutant. This might indicate that CP even dampens virulence through the WTA shielding effect.

Although the regulation of CP synthesis has been heavily studied [55], it is still unclear under which *in vivo* conditions CP is expressed. Interestingly, there is some evidence in murine infection models that even acapsular USA300 may produce CP under certain *in vivo* conditions [56]. CP expression on the skin of atopic dermatitis patients seems non-favorable for the bacterial population since strains with mutations within the *cap* operon are selected for on affected skin [22]. Interestingly, such mutations are not prevalent in isolates from acute infections. Increased capacity to adhere to skin structures might select for CP negative isolates in atopic dermatitis patients.

## Conclusion

CP expression masks WTA to protect the bacteria from opsonization with antibodies and infection by phages, which is likely beneficial for the bacteria during infection and/or colonization and to avoid phage attack e.g., during phage therapy. However, CP-dependent masking of bacterial adhesins is presumably a disadvantage for the bacteria. The highly heterogeneous, and tightly regulated CP expression in *S. aureus* has likely evolved as a bet hedging strategy to ensure that part of the population can fulfill important WTA-mediated functions like endothelial adhesion, while simultaneously evading detection by the host’s immune system or predatory phages.

## Ethics statement

Animal experiments were performed in strict accordance with the German regulations of the Society for Laboratory Animal Science (GV-SOLAS) and the European Health Law of the Federation of Laboratory Animal Science Associations (FELASA). The protocol was approved by the “Regierungspräsidium Tübingen” (permit no. IMIT 01/20G).

## Acknowledgements

We thank Cosima Hirt and Dorothee Kretschmer for assistance and scientific discussion regarding the animal experiments. We thank Simon Heilbronner for help with the ethics proposal and Libera Lo Presti for scientific discussions and editing of the manuscript.

## References

1. Chan, Y.G., et al., Staphylococcus aureus mutants lacking the LytR-CpsA-Psr family of enzymes release cell wall teichoic acids into the extracellular medium. J Bacteriol, 2013. 195(20): p. 4650–9.

2. Chan, Y.G., et al., The capsular polysaccharide of Staphylococcus aureus is attached to peptidoglycan by the LytR-CpsA-Psr (LCP) family of enzymes. J Biol Chem, 2014. 289(22): p. 15680–90.

3. Brown, S.S.M., John P. Jr.; Walker, Suzanne, Wall teichoic acids of gram-positive bacteria. Annual review of microbiology, 2013. 67: p. 313–336.

4. Weidenmaier, C. and J.C. Lee, Structure and Function of Surface Polysaccharides of Staphylococcus aureus. Curr Top Microbiol Immunol, 2016.

5. Weidenmaier, C. and A. Peschel, Teichoic acids and related cell-wall glycopolymers in Gram-positive physiology and host interactions. Nat Rev Microbiol, 2008. 6(4): p. 276–87.

6. Keinhörster, D., et al., Function and regulation of Staphylococcus aureus wall teichoic acids and capsular polysaccharides. Int J Med Microbiol, 2019: p. 151333.

7. Ingmer, H., D. Gerlach, and C. Wolz, Temperate Phages of Staphylococcus aureus. Microbiol Spectr, 2019. 7(5).

8. van Dalen, R.P. A.; van Sorge, N. M., Wall Teichoic Acid in Staphylococcus aureus Host Interaction. Trends Microbiol, 2020. 28(12): p. 985–998.

9. Wanner, S.S. J.; Keinhorster, D.; Weller, N.; George, S. E.; Kull, L.; Bauer, J.; Grau, T.; Winstel, V.; Stoy, H.; Kretschmer, D.; Kolata, J.; Wolz, C.; Broker, B. M.; Weidenmaier, C., Wall teichoic acids mediate increased virulence in Staphylococcus aureus. Nat Microbiol, 2017. 2: p. 16257.

10. Xia, G., et al., Wall teichoic Acid-dependent adsorption of staphylococcal siphovirus and myovirus. J Bacteriol, 2011. 193(15): p. 4006–9.

11. O’Riordan, K. and J.C. Lee, Staphylococcus aureus capsular polysaccharides. Clin Microbiol Rev, 2004. 17(1): p. 218–34.

12. Fournier, J.M.V., Willie. F.; and Karakawa, Walter. W., Purification and Characterization of Staphylococcus aureus Type 8 Capsular Polysaccharide. Infection and Immunity, 1984.

13. Moreau, M., et al., Structure of the type 5 capsular polysaccharide of Staphylococcus aureus. Carbohydr Res, 1990. 201(2): p. 285–97.

14. Jones, C., Revised structures for the capsular polysaccharides from Staphylococcus aureus Types 5 and 8, components of novel glycoconjugate vaccines. Carbohydr Res, 2005. 340(6): p. 1097–106.

15. Tuchscherr, L., et al., Staphylococcus aureus adaptation to the host and persistence: role of loss of capsular polysaccharide expression. Future Microbiol, 2010. 5(12): p. 1823–32.

16. Thakker, M., et al., Staphylococcus aureus serotype 5 capsular polysaccharide is antiphagocytic and enhances bacterial virulence in a murine bacteremia model. Infect Immun, 1998. 66(11): p. 5183–9.

17. Watts, A., et al., Staphylococcus aureus strains that express serotype 5 or serotype 8 capsular polysaccharides differ in virulence. Infect Immun, 2005. 73(6): p. 3502–11.

18. Nilsson, I.M., et al., The role of staphylococcal polysaccharide microcapsule expression in septicemia and septic arthritis. Infect Immun, 1997. 65(10): p. 4216–4221.

19. McLoughlin, R.M., et al., CD4+ T cells and CXC chemokines modulate the pathogenesis of Staphylococcus aureus wound infections. Proc Natl Acad Sci U S A, 2006. 103(27): p. 10408–10413.

20. Lattar, S.M., et al., Capsule expression and genotypic differences among Staphylococcus aureus isolates from patients with chronic or acute osteomyelitis. Infect Immun, 2009. 77(5): p. 1968–75.

21. Herbert, S., et al., Regulation of Staphylococcus aureus capsular polysaccharide type 5: CO2 inhibition in vitro and in vivo. J Infect Dis, 1997. 176(2): p. 431–438.

22. Key, F.M., et al., On-person adaptive evolution of Staphylococcus aureus during treatment for atopic dermatitis. Cell Host Microbe, 2023. 31(4): p. 593–603 e7.

23. Suligoy, C.M., et al., Acapsular Staphylococcus aureus with a non-functional agr regains capsule expression after passage through the bloodstream in a bacteremia mouse model. Sci Rep, 2020. 10(1): p. 14108.

24. Tuchscherr, L.P., et al., Capsule-negative Staphylococcus aureus induces chronic experimental mastitis in mice. Infect Immun, 2005. 73(12): p. 7932–7937.

25. Baddour, L.M., et al., Staphylococcus aureus microcapsule expression attenuates bacterial virulence in a rat model of experimental endocarditis. J Infect Dis, 1992. 165(4): p. 749–753.

26. Nemeth, J. and J.C. Lee, Antibodies to capsular polysaccharides are not protective against experimental Staphylococcus aureus endocarditis. Infect Immun, 1995. 63(2): p. 375–380.

27. Boyle-Vavra, S., et al., USA300 and USA500 clonal lineages of Staphylococcus aureus do not produce a capsular polysaccharide due to conserved mutations in the cap5 locus. mBio, 2015. 6(2).

28. Pöhlmann-Dietze, P., et al., Adherence of Staphylococcus aureus to endothelial cells: influence of capsular polysaccharide, global regulator agr, and bacterial growth phase. Infect Immun, 2000. 68(9): p. 4865–71.

29. Risley, A.L., et al., Capsular polysaccharide masks clumping factor A-mediated adherence of Staphylococcus aureus to fibrinogen and platelets. J Infect Dis, 2007. 196(6): p. 919–27.

30. Bae, T. and O. Schneewind, Allelic replacement in Staphylococcus aureus with inducible counter-selection. Plasmid, 2006. 55(1): p. 58–63.

31. Monk, I.R., et al., Transforming the untransformable: application of direct transformation to manipulate genetically Staphylococcus aureus and Staphylococcus epidermidis. MBio, 2012. 3(2).

32. Mainiero, M., et al., Differential target gene activation by the Staphylococcus aureus two-component system saeRS. J Bacteriol, 2010. 192(3): p. 613–23.

33. Peschel, A., et al., Inactivation of the dlt operon in Staphylococcus aureus confers sensitivity to defensins, protegrins, and other antimicrobial peptides. J Biol Chem, 1999. 274(13): p. 8405–10.

34. Chen, P.S., T.Y. Toribara, and H. Warner, Microdetermination of Phosphorus. Analytical Chemistry, 1965. 28, NO. 11.

35. Kropinski, A.M., et al., Enumeration of bacteriophages by double agar overlay plaque assay. Methods Mol Biol, 2009. 501: p. 69–76.

36. Hendriks, A., et al., Impact of Glycan Linkage to Staphylococcus aureus Wall Teichoic Acid on Langerin Recognition and Langerhans Cell Activation. ACS Infect Dis, 2021. 7(3): p. 624–635.

37. Driguez, P.-A., et al., Immunogenic Compositions Against S. aureus. 2017, Sanofi Pasteur: WO.

38. Lehar, S.M., et al., Novel antibody-antibiotic conjugate eliminates intracellular S. aureus. Nature, 2015. 527(7578): p. 323–8.

39. Di Carluccio, C., et al., Antibody Recognition of Different Staphylococcus aureus Wall Teichoic Acid Glycoforms. ACS Cent Sci, 2022. 8(10): p. 1383–1392.

40. Grundmeier, M., et al., Truncation of fibronectin-binding proteins in Staphylococcus aureus strain Newman leads to deficient adherence and host cell invasion due to loss of the cell wall anchor function. Infect Immun, 2004. 72(12): p. 7155–63.

41. van Dalen, R., A. Peschel, and N.M. van Sorge, Wall Teichoic Acid in Staphylococcus aureus Host Interaction. Trends Microbiol, 2020.

42. Baur, S., et al., A nasal epithelial receptor for Staphylococcus aureus WTA governs adhesion to epithelial cells and modulates nasal colonization. PLoS Pathog, 2014. 10(5): p. e1004089.

43. van Dalen, R., et al., Langerhans Cells Sense Staphylococcus aureus Wall Teichoic Acid through Langerin To Induce Inflammatory Responses. MBio, 2019. 10(3): p. e00330–19.

44. Weidenmaier, C., et al., Lack of wall teichoic acids in Staphylococcus aureus leads to reduced interactions with endothelial cells and to attenuated virulence in a rabbit model of endocarditis. J Infect Dis, 2005. 191(10): p. 1771–7.

45. Du, X., et al., Staphylococcus epidermidis clones express Staphylococcus aureus-type wall teichoic acid to shift from a commensal to pathogen lifestyle. Nat Microbiol, 2021. 6(6): p. 757–768.

46. Leprince, A. and J. Mahillon, Phage Adsorption to Gram-Positive Bacteria. Viruses, 2023. 15(1).

47. Bertozzi Silva, J., Z. Storms, and D. Sauvageau, Host receptors for bacteriophage adsorption. FEMS Microbiol Lett, 2016. 363(4).

48. Negus, D., et al., Poly-gamma-(D)-glutamic acid capsule interferes with lytic infection of Bacillus anthracis by B. anthracis-specific bacteriophages. Appl Environ Microbiol, 2013. 79(2): p. 714–7.

49. Wilkinson, B.J. and K.M. Holmes, Staphylococcus aureus cell surface: capsule as a barrier to bacteriophage adsorption. Infect Immun, 1979. 23(2): p. 549–52.

50. Sompolinsky, D., et al., Encapsulation and capsular types in isolates of Staphylococcus aureus from different sources and relationship to phage types. J Clin Microbiol, 1985. 22(5): p. 828–34.

51. Witte, W., Capsule formation in Staphylococcus aureus as a reason for nontypability by phages (author’s transl). Zentralblatt fur Bakteriologie, Parasitenkunde, Infektionskrankheiten und Hygiene. Erste Abteilung Originale. Reihe A: Medizinische Mikrobiologie und Parasitologie, 1975. 233(4): p. 447–451.

52. George, S.E., et al., Phenotypic heterogeneity and temporal expression of the capsular polysaccharide in Staphylococcus aureus. Mol Microbiol, 2015. 98(6): p. 1073–88.

53. Cunnion, K.M., J.C. Lee, and M.M. Frank, Capsule production and growth phase influence binding of complement to Staphylococcus aureus. Infect Immun, 2001. 69(11): p. 6796–803.

54. Portoles, M., et al., Staphylococcus aureus Cap5O has UDP-ManNAc dehydrogenase activity and is essential for capsule expression. Infect Immun, 2001. 69(2): p. 917–23.

55. Keinhörster, D., et al., Revisiting the regulation of the capsular polysaccharide biosynthesis gene cluster in Staphylococcus aureus. Mol Microbiol, 2019. 112(4): p. 1083–1099.

56. Mohamed, N., et al., Molecular epidemiology and expression of capsular polysaccharides in Staphylococcus aureus clinical isolates in the United States. PLoS One, 2019. 14(1): p. e0208356.

