## Supplemental Table 1 - 5 for "The capsular polysaccharide obstructs wall teichoic acid functions in *Staphylococcus aureus*"

### Supplemental tables

**Table S1. List of bacterial strains.**

| Bacterial strain | Description | Reference |
|---|---|---|
| ***S. aureus*** | | |
| USA300 JE2 | CA-MRSA, USA300 LAC cured of all 3 native plasmids, natural CP5 deficient | [1] |
| USA300 JE2 Cap | Mutations in CP-operon have been reversed, CP5 expression | [2] |
| USA300 JE2 Cap-high | Modification of Pcap SigB-binding-site, CP5 overexpression | [2] |
| USA300 JE2 Δ*tagO* | *tagO* deletion, WTA deficient mutant | Manuscript under revision |
| USA300 JE2 *srtA*::erm | Transposon insertion in *srtA*, mutant deficient for all major cell wall proteins | This work |
| Newman | HA-MRSA, CP5 | [3] |
| Newman high | Modification of Pcap SigB-binding-site, CP5 overexpression | [4] |
| Newman Δ*tagO* | *tagO* deletion, WTA deficient mutant | |
| RN4220 | Restriction-deficient derivate of 8325-4, indicator strain, phage propagation | [5] |
| LS1 | Mouse arthritis/osteomyelitis strain, Indicator strain for Φ13kΔ*int* | [6] |
| ***E. coli*** | | |
| DC10B | Cloning strain, DNA cytosine methyltransferase mutant | [7] |

**Table S2. List of bacteriophages.**

| Phages | Description | Reference |
|---|---|---|
| Φ11 | siphovirus, receptor: α-1,4-GlcNAc-, β-1,4-GlcNAc- and partially β-1,3-GlcNAc-RboP-WTA | |
| ΦK | myovirus, receptor: RboP-WTA | |
| Φ13k Δ*int* | siphoviurs, unknown phage receptor | This work |

**Table S3. List of plasmids and vectors**

| Plasmids | Description | Reference |
|---|---|---|
| pBASE6 | tetracyclin inducible suicide mutagenesis plasmid | [8] |
| pCG472 | pBASE6 for generation of USA300 JE2 Δ*tagO* | This work |

**Table S4. List of oligonucleotides.**

| Primer | extentionSEQUENCE | Use |
|---|---|---|
| tagOGibMutfor | ctcatcgcagtgcagcggAATAATGATA GCACATCATTTTGT | generation of USA300 JE2 Δ*tagO* |
| tagOGibMutrev | ccgggtaccgagctccggGCATTTATTG CTGCAATAAA | generation of USA300 JE2 Δ*tagO* |
| tagOlinkMutfor | aggtgaataaGGAATGAAAGCATAG CTGTATGGG | generation of USA300 JE2 Δ*tagO* |
| tagOlinkMutrev | ctttcattccTTATTCACCTTCATCGA TATT | generation of USA300 JE2 Δ*tagO* |
| tagocontrolfor | TCCTTTAATTGACCACTAGCT | control PCR |
| tagocontrolrev | AAGTACACGTTTATGGCAGT | control PCR |

**Table S5. List of eukaryotic cells.**

| Cell lines | Description | Company |
|---|---|---|
| HUVECs | primary Human Umbilical Vein Endothelial Cells | PromoCell (C-12203) |

### References

1. Nuxoll, A.S., et al., *CcpA Regulates Arginine Biosynthesis in Staphylococcus aureus through Repression of Proline Catabolism.* PLOS Pathogens, 2012. **8**(11): p. e1003033.
2. Keinhörster, D., et al., *Revisiting the regulation of the capsular polysaccharide biosynthesis gene cluster in Staphylococcus aureus.* Mol Microbiol, 2019. **112**(4): p. 1083-1099.
3. Duthie, E.S. and L.L. Lorenz, *Staphylococcal coagulase; mode of action and antigenicity.* J Gen Microbiol, 1952. **6**(1-2): p. 95-107.
4. Keinhörster, D., et al., *Function and regulation of Staphylococcus aureus wall teichoic acids and capsular polysaccharides.* Int J Med Microbiol, 2019: p. 151333.

5. Kreiswirth, B.N., et al., *The toxic shock syndrome exotoxin structural gene is not detectably transmitted by a prophage.* Nature, 1983. **305**(5936): p. 709-12.
6. Bremell, T., et al., *Outbreak of spontaneous staphylococcal arthritis and osteitis in mice.* Arthritis Rheum, 1990. **33**(11): p. 1739-44.
7. Monk, I.R., et al., *Transforming the untransformable: application of direct transformation to manipulate genetically Staphylococcus aureus and Staphylococcus epidermidis.* MBio, 2012. **3**(2).
8. Geiger, T., et al., *The stringent response of Staphylococcus aureus and its impact on survival after phagocytosis through the induction of intracellular PSMs expression.* PLoS Pathog, 2012. **8**(11): p. e1003016.
